# Dosimetric and biologic intercomparison between electron and proton FLASH beams

**DOI:** 10.1101/2023.04.20.537497

**Authors:** A Almeida, M Togno, P Ballesteros-Zebadua, J Franco-Perez, R Geyer, R Schaefer, B Petit, V Grilj, D Meer, S Safai, T Lomax, DC Weber, C Bailat, S Psoroulas, MC Vozenin

## Abstract

**Background and purpose:** The FLASH effect has been validated in different preclinical experiments with electrons (eFLASH) and protons (pFLASH) operating at a mean dose rate above 40 Gy/s. However, no systematic intercomparison of the FLASH effect produced by e *vs*. pFLASH has yet been performed and constitutes the aim of the present study.

**Materials and methods:** The electron eRT6/Oriatron/CHUV/5.5 MeV and proton Gantry1/PSI/170 MeV were used to deliver conventional (0.1 Gy/s eCONV and pCONV) and FLASH (≥100 Gy/s eFLASH and pFLASH) irradiation. Protons were delivered in transmission. Dosimetric and biologic intercomparisons were performed with previously validated models.

**Results:** Doses measured at Gantry1 were in agreement (± 2.5%) with reference dosimeters calibrated at CHUV/IRA. The neurocognitive capacity of e and pFLASH irradiated mice was indistinguishable from the control while both e and pCONV irradiated cohorts showed cognitive decrements. Complete tumor response was obtained with the two beams and was similar between e and pFLASH *vs*. e and pCONV. Tumor rejection was similar indicating that T-cell memory response is beam-type and dose-rate independent.

**Conclusion:** Despite major differences in the temporal microstructure, this study shows that dosimetric standards can be established. The sparing of brain function and tumor control produced by the two beams were similar, suggesting that the most important physical parameter driving the FLASH effect is the overall time of exposure which should be in the range of hundreds of milliseconds for WBI in mice. In addition, we observed that immunological memory response is similar between electron and proton beams and is independent off the dose rate.

## Introduction

Ultra-high dose rate FLASH radiotherapy (FLASH) rapidly became a new field of research and investigations thanks to its ability to open the therapeutic window by enhancing the differential response between normal tissue and tumors. For a given dose, FLASH preserves normal tissue in various species and organs while simultaneously maintaining the anti-tumor efficacy equivalent to radiotherapy delivered at a conventional dose rate (CONV). This beneficial biological outcome has been names the “FLASH effect” *(1)*.

So far, most of the experimental studies available were performed with a dedicated Oriatron/eRT6 experimental electron beam located at the CHUV. However, the low energy (5.5 MeV) of this electron irradiator *(2)* and similar electron devices *(3, 4)* now available on the market limits the applicability of FLASH to small animals and/or superficial tumors. Therefore, investigating the FLASH effect on alternative beams is required for efficient clinical translation. Recently, clinical proton beams have been optimized to operate at a mean dose rate above 40 Gy/s *(5)* and qualitatively similar findings showing normal tissue sparing and sustained anti-tumor response have been validated in preclinical experiments using rodent gut *(5, 6)* and skin *(7)* models. Normal tissue sparing was also described using different modalities of proton-FLASH-PBS *(8)* and Bragg peak proton-FLASH beams *(9, 10)*. Although the temporal structures of e and pFLASH beams are different, both have been shown to produce the FLASH effect in mice.

Subtle differences might still be relevant as shown in a recent study conducted in our group where physico-chemical and biological differences between e and pFLASH beams have been observed in zebrafish (ZF) embryos *(11)*. In addition and despite many positive findings, it is important to note that other studies have reported the absence of the FLASH effect using proton *(12)* and electron *(13)* beams and two recent clinical studies have reported no differences between FLASH and CONV delivered with electron *(14)* and proton FLASH beams *(15)*. Although performed at a high-dose rate, these studies might have been carried out with suboptimal conditions and/or beyond those required to elicit a significant FLASH effect. In addition, a recent veterinary phase III clinical trial showed that a high single dose could cause severe late toxicity even when irradiation using a FLASH validated beam is used *(16)*. These negative results emphasize the need to carefully document and control experimental conditions (physical and biological parameters) required to produce the FLASH effect. Standardization of parameters and conditions will allow for reproducible and comparable experimental results that will ultimately promote the safe and reliable transfer of these strategies to the clinic.

Consequently, this study has been designed to compare electrons and protons delivered at FLASH and CONV dose rates, both dosimetrically and biologically. First, we show that a dosimetric consensus strategy recently developed for eFLASH *(17)* is applicable to proton beams. Second, we used the model of late tissue damages developed for eFLASH *(18–20)* to validate the capability of protons to achieve pFLASH sparing effect at 110 Gy/s. Finally, anti-tumor efficacy was confirmed to be dose rate and beam type independent, as was the induction of *in situ* vaccination. This study validates PSI Gantry1 as a proton beam capable of generating a FLASH effect and shows that dosimetric and biologic consensus can be standardize across various facilities. And radiation qualities

## Material and Method

### Irradiation devices

Irradiation was performed using

1. The Oriatron 6e (eRT6; PMB-Alcen, Peynier, France), a 5,5 MeV electron beam linear accelerator (LINAC) described previously *(21)* and extensively validated to produce the FLASH effect *(1)*. The prototype was operated at 0.1 Gy/s for CONV and at ≥100Gy/s for FLASH. The beam parameters used in this study are shown in Table 1.

**Table 1.** 2. The PSI Comet Cyclotron delivers a 250MeV proton beam, which is transported with ∼85% efficiency to Gantry1 as already described *(22)*. The beam was degraded to 170 MeV at the nozzle exit and delivered 0.1Gy/s for CONV, and 110Gy/s for FLASH in transmission mode. The beam parameters used in this study are included in Table 2.

| Mode | Prescribed Dose (Gy) | Dose Rate (Gy/s) | Beam parameters |  |  |  |  |
| --- | --- | --- | --- | --- | --- | --- | --- |
| | | | Frequency (Hz) | SSD (mm) | Pulse width (μs) | Number of pulses | Treatment time (s) |
| eCONV | 10 | 0.1 | 10 | 800 | 1 | >600 | >60 |
|  | 20 | 0.1 | 10 | 612 | 1 | >1500 | >120 |
| eFLASH | 10 | $5.5 \cdot 10^6$ | na | 350 | 1.8 | 1 | $1.8 \cdot 10^{-6}$ |
|  | 10 | 100 | 100 | 920 | 1.8 | 10 | 0.09 |
|  | 20 | 2000 | 100 | 320 | 1.8 | 2 | 0.01 |

**Table 2.**

| Mode | Prescribed Dose (Gy) | Dose Rate (Gy/s) | Beam parameters |  |  |
| --- | --- | --- | --- | --- | --- |
|  |  |  | Frequency (Hz) | Cyclotron beam current (nA) | Treatment time (s) |
| pCONV | 10 | 0.1 | $7.285 \cdot 10^7$ | 0,66 | 100 |
| | 20 | 0.1 | $7.285 \cdot 10^7$ | 0.66 | 200 |
| pFLASH | 10 | 110 | $7.285 \cdot 10^7$ | 750 | 0.09 |
| | 20 | 110 | $7.285 \cdot 10^7$ | 750 | 0.18 |

### Dosimetric intercomparison

A cuboid phantom (25 × 25 × 32 mm^3^) made of acrylic (PMMA, ρ = 1.19 g·cm^-3^) and recently described and validated by Jorge et al., was used for this comparison *(23)*. The phantom has a 5 mm (diameter) by 10.4 mm (length) central cylindrical cavity to simultaneously house three TLDs, two alanine pellets, and six laser-cut films. Irradiations were performed in the same conditions used for the biological comparison: a graphite applicator (13.0 × 13.0 × 2.5 cm^3^) with 17 mm^2^ diameter aperture was used at CHUV-eRT6, while a 6 cm thick copper collimator with 17 mm^2^ diameter aperture was used at PSI-Gantry1. We report additional details on the irradiation field and setup in the section 1.1 and 1.2 of the supplementary material.

Five PMMA phantoms were mailed to PSI and four of them were irradiated behind the collimator, with the rectangular face orthogonal to the main beam axis. The non-irradiated phantom served as a background monitor. After irradiation, the phantoms were sent back to IRA for readout. Dosimetry performed at CHUV-eRT6 has been detailed extensively in previous works *(2, 24)*. The standard uncertainty on the absorbed dose measurements using the passive dosimeters in the cuboid phantoms was obtained by quadratic summation of the contribution coming from the calibration factors and the reproducibility of the measurements in the non-reference geometry. We evaluated the standard uncertainty (k=1) at 4 % for TLDs, 3 % for alanine, and 4 % for the laser-cut films.

At PSI-Gantry1, the reference dosimetry is performed with EBT3 Gafchromic films (Ashland, Bridgewater, US) and a synthetic single-crystal microDiamond (PTW, Freiburg, Germany). In conditions using CONV, both detectors are cross-calibrated to a reference chamber traceable to the Swiss primary standard laboratory METAS. The detector responses at different dose rates were previously characterized by Togno et al.*(25)* and corrections to detectors reading were considered accordingly.

The overall combined uncertainty on absorbed dose to water in proton beams was 2.5 % (k=1) and 1.9 % (k=1) for EBT3 films and PTW microDiamond, respectively. This estimated uncertainty applies to both FLASH and CONV conditions. EBT3 films as well as microDiamond were used for comparison. In preparation for the mice experiments, the field dose was also measured with an Advanced Markus ion chamber (PTW, Freiburg, Germany) calibrated at METAS, with an overall uncertainty of 2.2 % (k=1).

### Biologic intercomparison

To enable biological comparison, irradiation settings were defined to besimilar between the two beams. The prescription dose for mice irradiations was determined by surface dose measurements on a 30 × 30 cm^2^ solid water slab positioned behind a 17 mm^2^ in diameter aperture of a graphite applicator (13.0 × 13.0 × 2.5 cm^3^). Animal experiments were approved by the Swiss (VD3603) ethics Committees for Animal Experimentation and performed within institutional guidelines.

### Normal brain response

#### Whole brain irradiations (WBIs)

Female C57BL/6J mice (n=10-12 animals per group) were purchased from CRL at the age of eight weeks. WBIs were performed under isoflurane anaesthesia. The mouse head was positioned behind and in contact with the 17 mm^2^ Ø applicator to irradiate the whole encephalon region while limiting the dose to the eyes, the mouth, and the rest of the body.

#### Novel object recognition testing

Neurocognitive impairments are typically found after treatments with CONV irradiation.

To determine the effects of FLASH and CONV using electron and proton on cognitive function, tumor-free animals were used to avoid confounding factors caused by tumor growth. Mice were irradiated whole brain with a single dose of 10 Gy delivered using FLASH (≥100Gy/s) or CONV (0.1 Gy/s) with eRT6 or Gantry1 parameters described in table 1 and 2. Novel Object Recognition (NOR) studies were performed 2 months post-RT, which is a time when alteration of hippocampal and frontal cortical learning and memory are stabilized. The NOR task was performed as previously described to validate the FLASH sparing effect *(18)*. It involved a sequence of habituation (no objects), familiarization (2 identical objects) and a final test phase in which one of the prior objects is switched with a different one. Animals tend to explore the novel object, and successful performance on this task relies on intact perirhinal cortex function *(26)*.

### Tumor response

#### Primary tumor irradiations and follow up

Female C57BL/6J mice (n=4-6 animals per group) were purchased from CRL at the age of eight weeks and used for subcutaneous implantation with 5 million murine GBM GL261 cells (Seligman, 1939) in the left flank. When tumor volume reached 80-100 mm^3^ they were locally irradiated with a single dose of 20 Gy using the 17 mm^2^ Ø applicator by stretching the skin and tumor over the applicator. Tumor growth was monitored by caliper measurement three times a week, and the volume was calculated with the formula of an oblate ellipsoid: V = (a × b^2^)/2, where <u>a</u> and <u>b</u> are the minor and major axes of the tumors.

#### Tumor rechallenge

GL261 cells are known to be highly aggressive but moderately immunogenic and radiosensitive *in vitro as* 2 Gy is already sufficient to generate 50 % of cell death *(27)*. To evaluate the potential of FLASH to generate *in situ* vaccine and T cell memory response, animals with a stable complete response for over 140 days post-RT were rechallenged with 5 × 10^6^ cells implanted in the opposite right flank. Tumor growth was monitored by caliper measurement.

#### Statistical analysis

Statistical analyses were performed using GraphPad Prism (version 9.1).

The normality of groups was assessed using the Shapiro-Wilk test. For NOR evaluation one-way ANOVA was used to determine the significance between all groups using Tuckey’s multiple comparisons test. For tumor response, P values were estimated from Kruskal-Wallis test using Dunn’s multiple comparison test. Results were expressed as mean ± SEM. All analyses considered a value of *P* ≤ 0.05 to be statistically significant.

## Results

### Dose measurements

Doses measured in preparation of the biological experiments are shown in **Figure 1**. The detectors were irradiated sequentially, hence the readings were corrected for beam output fluctuations (< 1.5%) between different deliveries at different dose rates. The dose measured with the three dosimeters agrees well within the experimental uncertainties, for both FLASH and CONV. The relative bias, i.e. the percentage difference between measured and expected dose, is in the range (−1.4 – 0) % for all detectors and dose rates.

**Figure 1.**
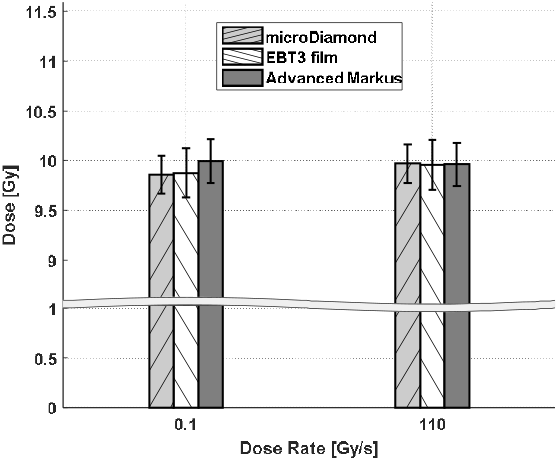
Results of dose measurements (target dose 10 Gy) using PSI dosimeters, at dose rates of 0.1 Gy/s and 110 Gy/s.

The results of the dosimetric comparison are shown in **Figure 2**. Two PMMA phantoms were irradiated using FLASH and CONV. **Figure 2A** shows, for the detectors loaded into the PMMA phantoms, the Co-60 reference values of absorbed dose to water provided by IRA. Thus, the values represent the absorbed dose to water that should be delivered in a Co-60 calibration beam to induce the same signal as measured in the Gantry1 proton beam. To provide consistency with the dose measured with PSI detectors, beam quality correction factors 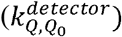 are necessary. Using the same approach as Palmans et al.*(28)*, we experimentally determined beam quality correction factors by cross-calibration against a reference ion-chamber in a proton beam at PSI Gantry2. The details are included in section 1.3 of the supplementary material –. The estimated correction factors are 1.00, 1.12 and 0.98 for Alanine, TLDs and Gafchromic films, respectively. The uncertainties are in the (2.3 – 5.5) % range with the uncertainty of the TLDs being significantly affected by measurement reproducibility. After correction with experimental 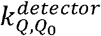 factors, the dose measured with Alanine and TLD detectors agreed within the standard uncertainty (k=1) with the dose measured at PSI with Gafchromic films and a microdiamond detector (**Figure 2B**). Laser-cut Gafchromic films in the PMMA phantoms slightly underestimated the dose with respect to the other detectors, but was consistent with Jorge et al.. The relative bias between the average dose measured with the PSI detector and the average dose measured in the audit phantoms was -1.9 % (0.1 Gy/s) and +2.5 % (110 Gy/s).

**Figure 2.**
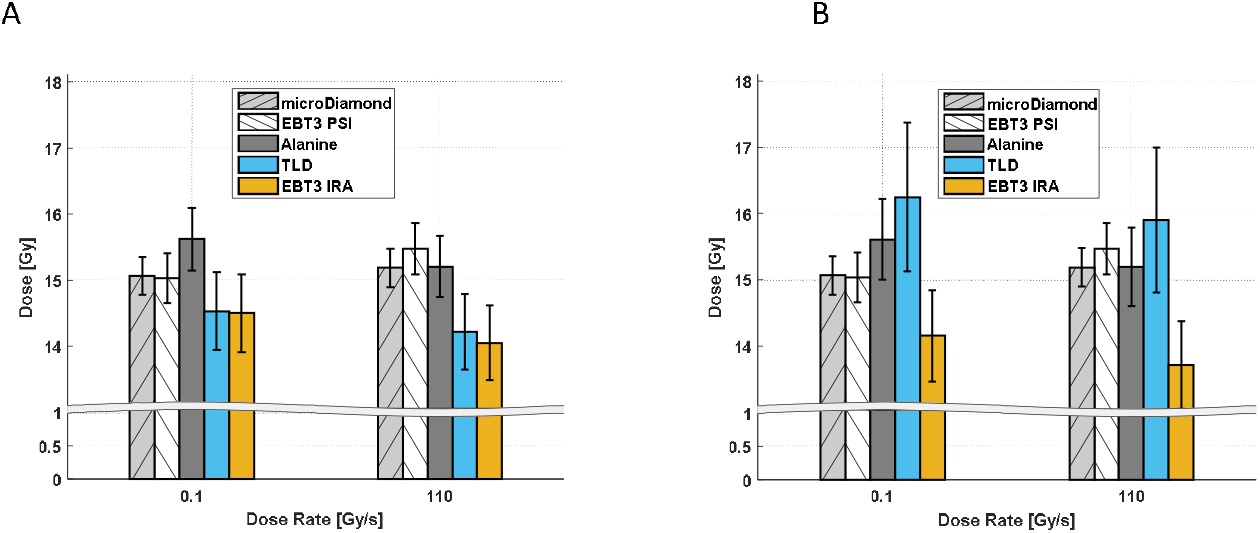
Results of the dosimetric intercomparison between PSI and IRA dosimeters, at dose rates of 0.1 Gy/s and 110 Gy/s. The dose measured with IRA dosimeters (Alanine, TLD, EBT3 IRA) is reported as Co-60 absorbed dose to water (A) and with experimentally determined corrections for beam quality applied (B). Error bars represent the combined standard uncertainty (k=1).

### Absence of cognitive impairment is validated with pFLASH

Animals (n=10-12) exposed to WBI with 10 Gy e and pFLASH and subjected to the NOR task (**Figure 3**) were statistically indistinguishable from controls, whereas CONV irradiated cohorts exhibited a reduction in their recognition ratio. Cognitive protection after eFLASH using either 10^7^Gy/s or 100 Gy/s were similar using the eRT6 electron beam. Importantly, mice from all cohorts which were subjected to spontaneous exploration tasks exhibited normal motor function and exploration.

**Figure 3.**
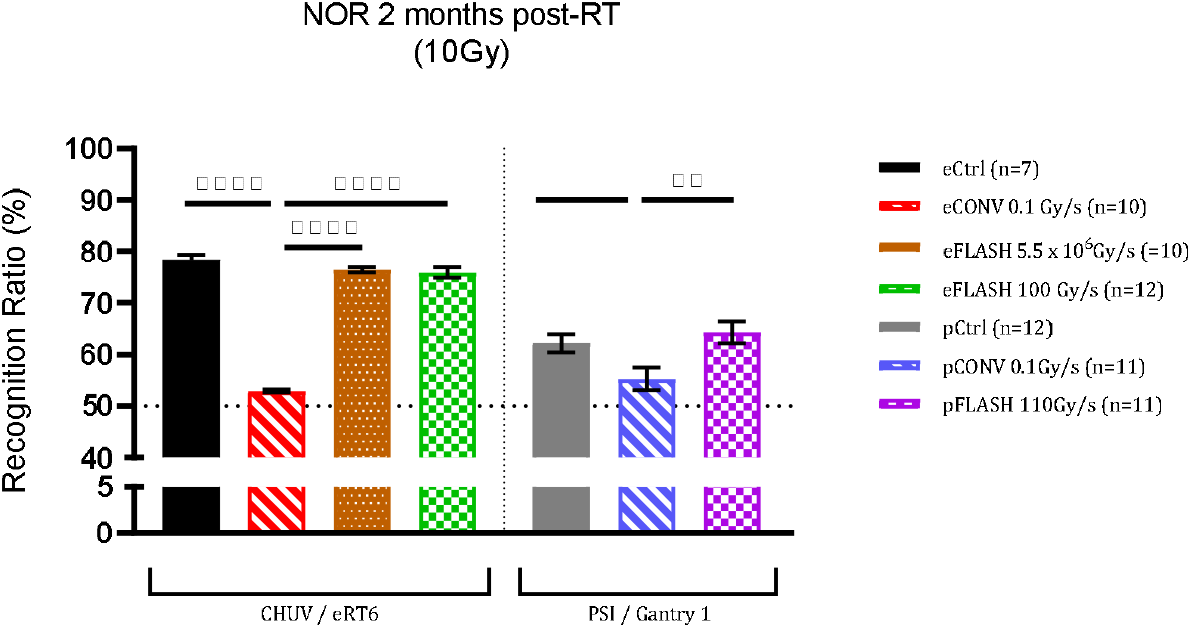
Novel Object Recognition Test: Animals exposed to 10 Gy with both eFLASH and pFLASH, have statistically indistinguishable recognition ratios relative to controls indicating a preference for the novel object, whereas mice irradiated with eCONV and pCONV showed impairment compared to controls. Mean ± SEM (n = 10–12 per group); p-values were compared against CONV and derived from One-way ANOVA followed by Tukey’s correction for multiple comparisons. * p < 0.05, **p < 0.01**p < 0.001.

### Complete anti-tumor response is beam-type and dose-rate independent

Five groups of C57BL/6J mice (n=4-6) subcutaneously implanted with GL261 murine GBM model were irradiated (or not control) with single doses of 20 Gy FLASH and CONV using both electron (CHUV/eRT6) and proton (PSI/Gantry1) beams. All tumor-bearing animals showed a complete and long-term response after irradiation, irrespective of the beam and dose rate used (**Figure 4A and C**). No tumor relapse occurred in the irradiated cohorts >140 days post-irradiation.

**Figure 4.**
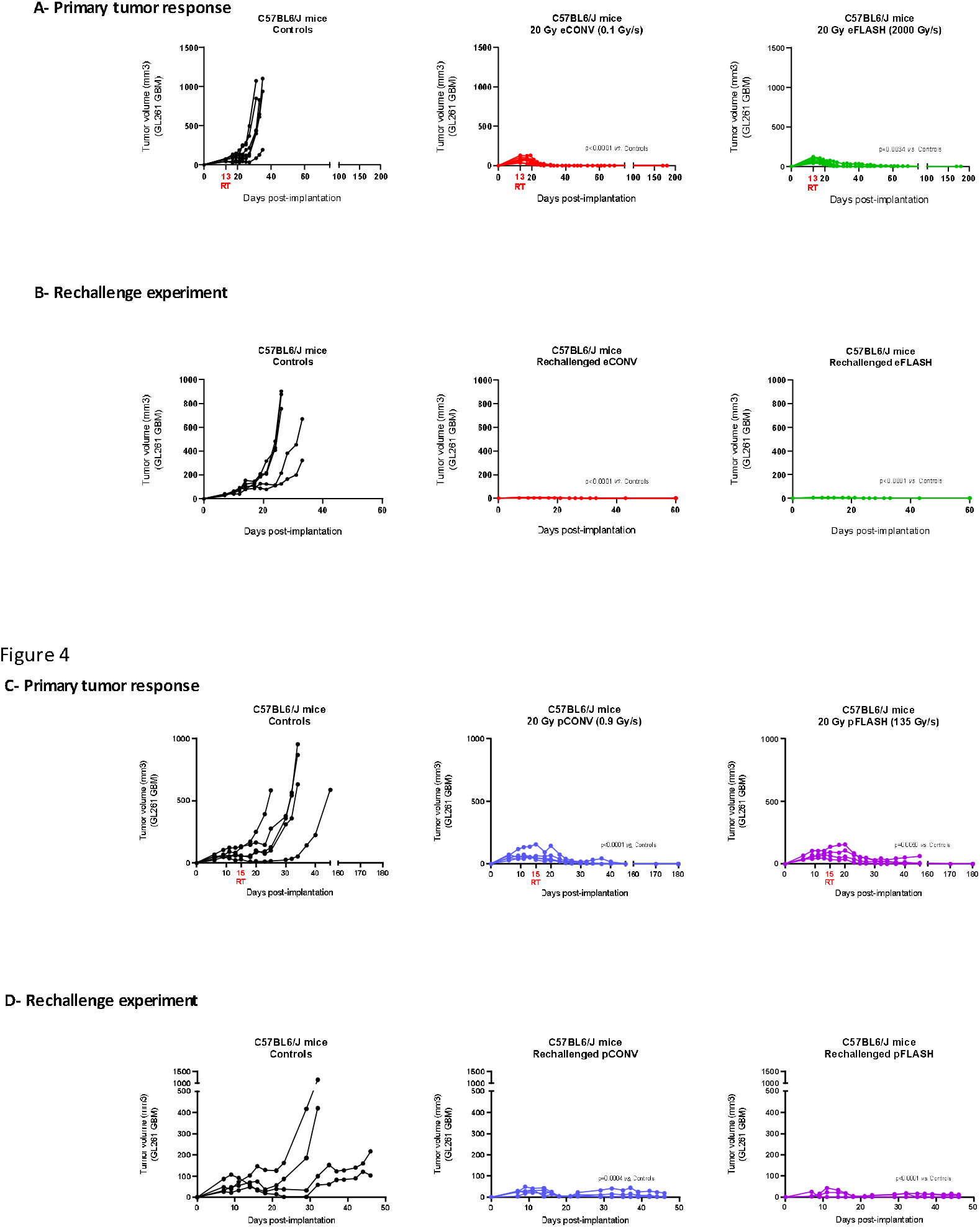
e/pFLASH and e/pCONV are equipotent in curing animals and generate a similar immunological memory response against GL261 cell line. GL261 glioblastoma (GBM) tumors were irradiated at 20 Gy with e/pFLASH or e/pCONV after subcutaneous engraftment into immunocompetent C57BL/6J female mice (A and C). Cured immunocompetent C57BL/6J female mice were rechallenged with 5 × 10^6^ cells implanted in the opposite flank (B and D). Tumor growth delay was followed by caliper measurement 3 times per week. Results are given in individual values. Statistical analysis of tumor growth curves was performed using Mann-Whitney test. ns (0.12), * (0.0332), ** (0.0021), *** (0.0002), **** (<0.0001).

### Radiation-induced in situ vaccination is beam-type and dose-rate independent

Since a long-term cure was achieved in all irradiated animals, we also evaluated the possible occurrence of a radiation-induced memory response. 140 days after the complete response **(Figure 4B and D)**, mice were rechallenged with GBM GL261 tumors engrafted on the opposite flank. While tumor growth occurred in 100% naïve control animals, rechallenged cohorts rejected tumors supporting the notion that radiation-induced T cell memory response was long-lasting, beam type and dose rate independent.

## Discussion

This paper is the first to report a systematic comparison of electron and proton FLASH capability and the first successful report of the FLASH effect with the Gantry1 proton beam using ultra high dose rate irradiation. This study shows that despite major differences in the temporal structure between electron and proton beams, dosimetric standards can be established and used for subsequent radiobiological evaluation investigating the FLASH effect. At the biological level, electron and proton beams delivering FLASH induced similar long-term benefit of neurocognitive sparing as well as complete tumor response associated with sustained T memory response. This study suggests that the definition of a common molecular pattern after e and pFLASH may be a key determinant of the FLASH effect.

Accurate dose determination is fundamental to conduct rigorous research on the FLASH effect, but it remains challenging at ultra-high dose rate beams. At PSI Gantry1, different detectors have been recently tested up to several kGy/s *(29, 30)*. To prepare for the biological studies, we verified that the measured field dose was reproducible and consistent between EBT3 Gafchromic films, Advanced Markus ion-chambers and synthetic microDiamonds, after proper characterization of the detectors *(25)*.

At present, no primary standard is available for proton beams, at either CONV or FLASH dose rates. In our study, we have extended a comparison scheme previously tested with MeV electron beams *(17)* to establish dosimetric consensus and cross-validation of Gantry1 and eRT6 irradiation beams. To provide consistency between the dose reported by PSI and IRA, beam quality correction factors were experimentally determined by cross-calibration of IRA detectors in proton beams. Although the reproducibility of the measurements varied in the (0.9 – 3.8) % range for different detectors, the measured correction factors for Alanine (1.00±0.02), TLDs (1.12±0.06) and EBT3 Gafchromic films (0.98±0.02) were found to be compatible with values reported in the literature *(28, 31–33)*. Clearly, a more precise determination of these correction factors is needed to further improve the accuracy of the measured dose to water in proton beams. Considering the agreement of all tested detectors within standard uncertainty (k=2), we successfully proved the dosimetric consistency of PSI and IRA/CHUV. Additionally, we showed that the methodology developed by Jorge et al. could be extended to proton beams *(23)*.

Most of the recent publications reporting the FLASH capabilities of novel beams have been conducted using GI, an acute responding organ *(6, 34)*. However, late normal tissue toxicity has always been and remains the main concern in the field of radiation oncology, as highlighted by our recent clinical study in domestic cat patients. In the latter study, acute toxicity was minor, whereas late osteoradionecrosis occurred in 3/7 cats irradiated with a single dose of 30 Gy with eRT6 (3 pulses, 1500 Gy/s) *(16)*. We therefore chose to validate the FLASH capability of Gantry1 protons using the brain as a model of late-responding tissue. For this, Gantry1 was operated at its maximal dose rate to achieve homogeneous dose and dose-rate coverage of the brain at 110 Gy/s. This dose rate was also used at eRT6 (100 Gy/s). Interestingly, the results of the NOR tests obtained at Gantry1 and eRT6 were similar 2 months post-RT showing cognitive sparing when a dose rate in excess of 100 Gy/s is used. Cognitive scores varied between the two experimental groups (electron and proton), this is an inerrant limitation of NOR as reviewed in Drayson et al., *(26)*, however these relative results are consistent with our previous dose rate escalation experiments *(18)* and support the idea that the FLASH-sparing effect can be reproduced in the brain of mice above 100 Gy/s independently of the beam used. More studies are ongoing to define a dose-modifying factor at Gantry1; however, our results suggest that the most important physical parameter to produce the FLASH-sparing effect is the overall time of irradiation exposure and that exposure in the range of hundreds of milliseconds is sufficient, at least when small volumes (mouse brain) are irradiated with a dose of 10 Gy.

In addition, while our recent study failed to produce a FLASH-sparing effect in ZF embryos *(11)*, the present study shows that Gantry1 is able to generate a FLASH-sparing effect in mice, using neurocognition as a functional outcome. Kacem et al. showed that ZF morphogenesis was protected by the proton beam after both FLASH and CONV, whereas the electron beam was damaging during CONV. These studies suggest subtle biological differences triggered by proton and electron beams depending on the biological target. They suggest that ZF embryos are more sensitive to the nature of the beam than mice and might require specific conditions as already described *(12, 35)*, and/or a higher dose rate (above 10^7^ Gy/s) to reveal the FLASH-sparing effect as observed in mouse tissues (around 100 Gy/s).

At the tumor level, the present study shows that the curative potential of irradiation is not modified by FLASH and also shows that long-term T cell memory response is activated by irradiation. However, we found no difference between electron and proton beams nor between FLASH and CONV. These results are consistent with a previous report *(36)* and are confirmed with pFLASH beams. Along with the numerous reports showing that tumor growth delay is similar between FLASH and CONV, the present study supports the idea that, unlike normal tissues, tumor response to ionizing radiation is dose rate independent. These results do not support any occurrence of a FLASH-specific immune response in tumors, as also reported in a recent study using an orthotopic glioma rat model *(37)*.

In summary, this study provides a strategy to validate new FLASH beams from the dosimetric to the biological endpoints. It also suggests that a dose rate in the range of 100 Gy/s delivered in less than a hundred milliseconds will be sufficient to produce the FLASH effect in a small volume. This information might be useful for clinical development of FLASH if this dose rate remains valid in humans and larger volumes.

## Acknowledgment

Funding was provided by National Institutes of Health grant P01CA244091-01 (to MCV supporting AA, VG); Swiss National Science Foundation grant Spirit IZSTZ0_198747/1 (to MCV and PBZ supporting JFC) and MAGIC -FNS CRS II5_186369 (to MCV and supporting VG); and CONACYT for supporting PBZ sabbatical in Switzerland. We also thank the Lausanne core facilities including the Animal facility, In vivo imaging facility, Mouse pathology facility at Epalinges.

## Supplementary material

### 1.1 Animal irradiation setup at PSI-Gantry1

At PSI-Gantry1, a 250 MeV proton beam is transported from the cyclotron to the treatment nozzle with an efficiency of ∼85%. In the nozzle, the pristine beam traverses 40 Polystyrene (PS) plates resulting in a residual energy of 170 MeV and a beam sigma of ∼ 25 mm at 1 m distance downstream of the nozzle. A 6 cm thick copper collimator is used to shape the 17 mm large irradiation field. Additional shielding is placed adjacent to the collimator to protect the animals from any stray radiation. An illustrative view of the irradiation setup is provided in Figure 1.

**Supplementary Figure 1.**
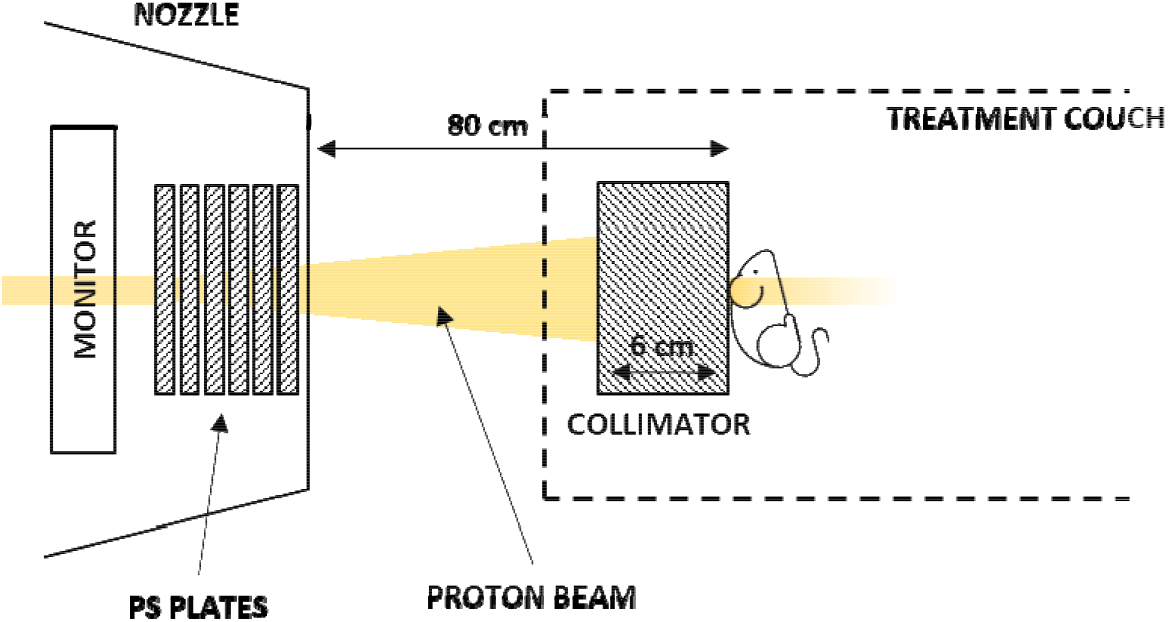
Illustrative representation of the animal irradiation setup at PSI Gantry1 – view from the top (dimensions are not to scale).

The dose delivery is supervised by a monitor chamber *(22)* built in the Gantry nozzle. On the day of the experiment, the output of the machine is verified and eventually corrected by means of a commercial synthetic microDiamond detector (PTW, Freiburg, Germany) at the required dose rates. Additionally, a TM7862 transmission chamber (PTW, Freiburg, Germany) is used as a redundant monitor. The response of the TM7862 ion chamber is typically characterized beforehand against a Faraday cup *(25)*.

### 1.2 Phantom irradiation setup at PSI-Gantry1

Two of the PMMA phantoms loaded with passive detectors were irradiated at each dose rate, i.e. conventional (0.1 Gy/s) and FLASH (110 Gy/s) dose rate. The phantoms were positioned downstream of the copper collimator, aligned using a wall-mounted laser, with the rectangular face orthogonal to the beam axis (Figure 2). The same irradiation setup was replicated for the measurements with PSI reference dosimeters. The following elements were placed downstream of the collimator, in sequence and attached to each other: 11 mm thick PMMA slab, EBT3 film, microDiamond detector (Figure 3). The microDiamond was inserted into a plastic phantom to ease the positioning with respect to the collimator. Note that the microDiamond has an intrinsic water equivalent buildup of 1.1 mm *(38)* and a sensitive circular area with a 2.2 mm diameter.

**Supplementary Figure 2.**
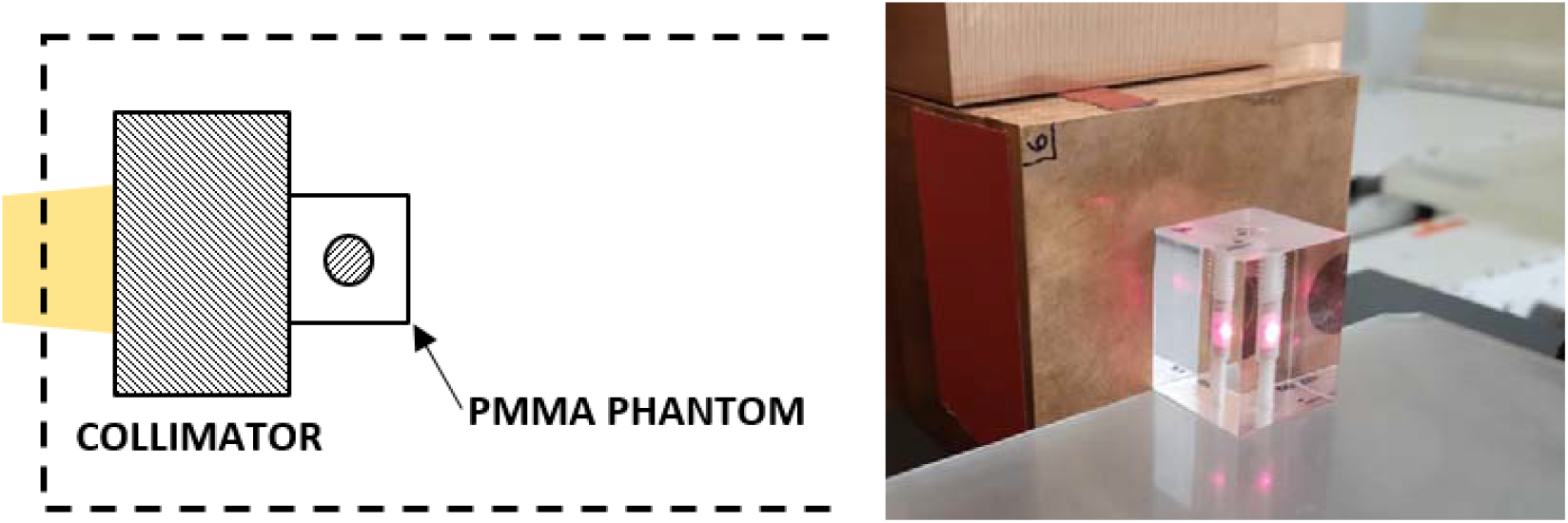
Schematic diagram and picture of the irradiation setup for the PMMA phantoms.

**Supplementary Figure 3.**
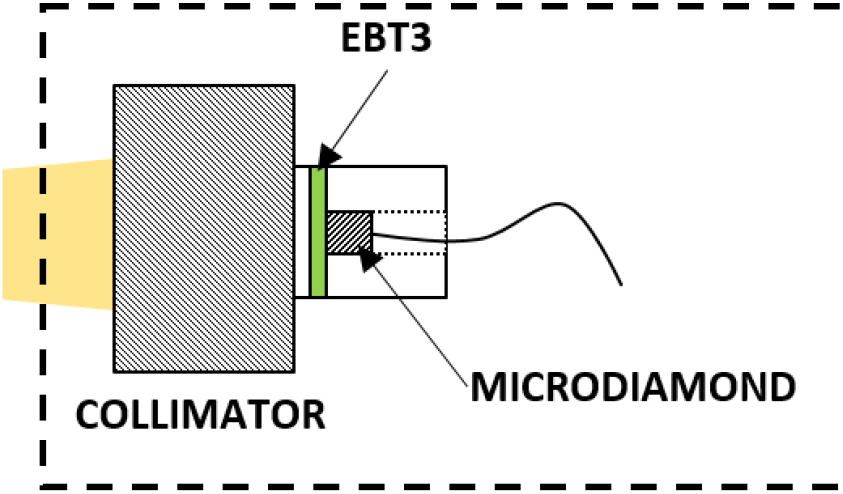
Schematic diagram of the irradaition setup for EBT3 films and microDiamond.

The 2D dose distributions measured with EBT3 films – for the setup illustrated in Figure 3 – are shown in Figures 4 and 5 for conventional and ultra-high dose rate, respectively. Orthogonal beam profiles are shown as well.

The EBT3 dose reported in the Results section is the average dose – at the center of the field – evaluated over an area equal to the microDiamond sensitive area.

**Supplementary Figure 4.**
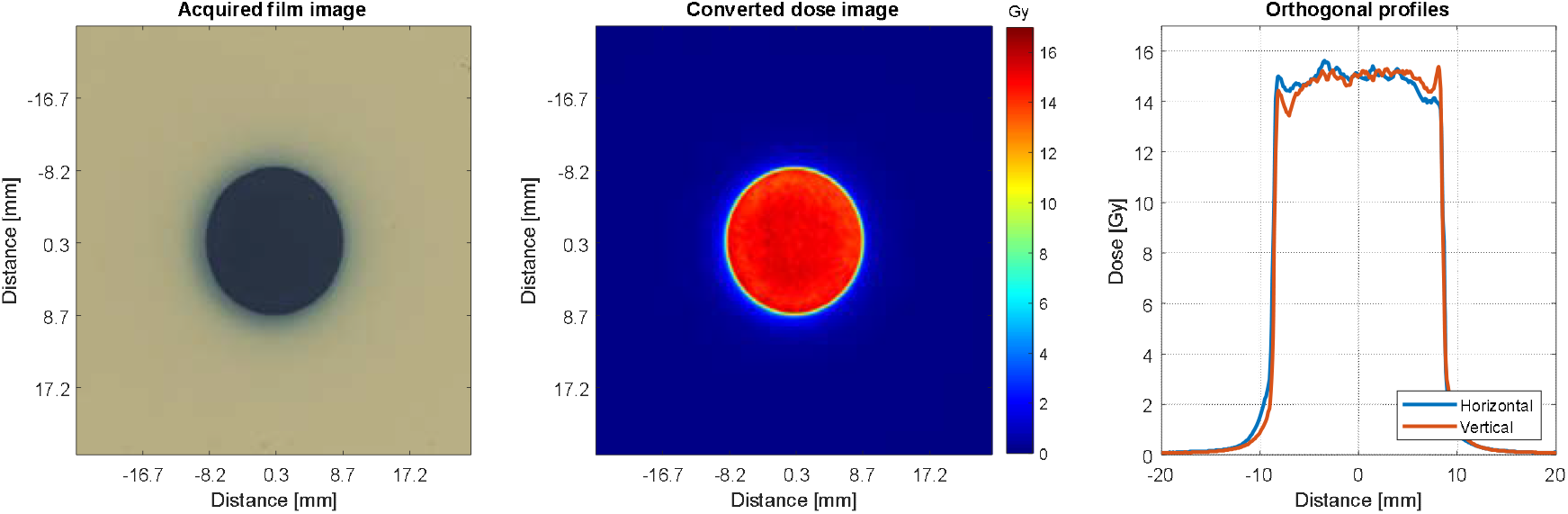
EBT3 measured dose at conventional dose rate (0.1 Gy/s).

**Supplementary Figure 5.**
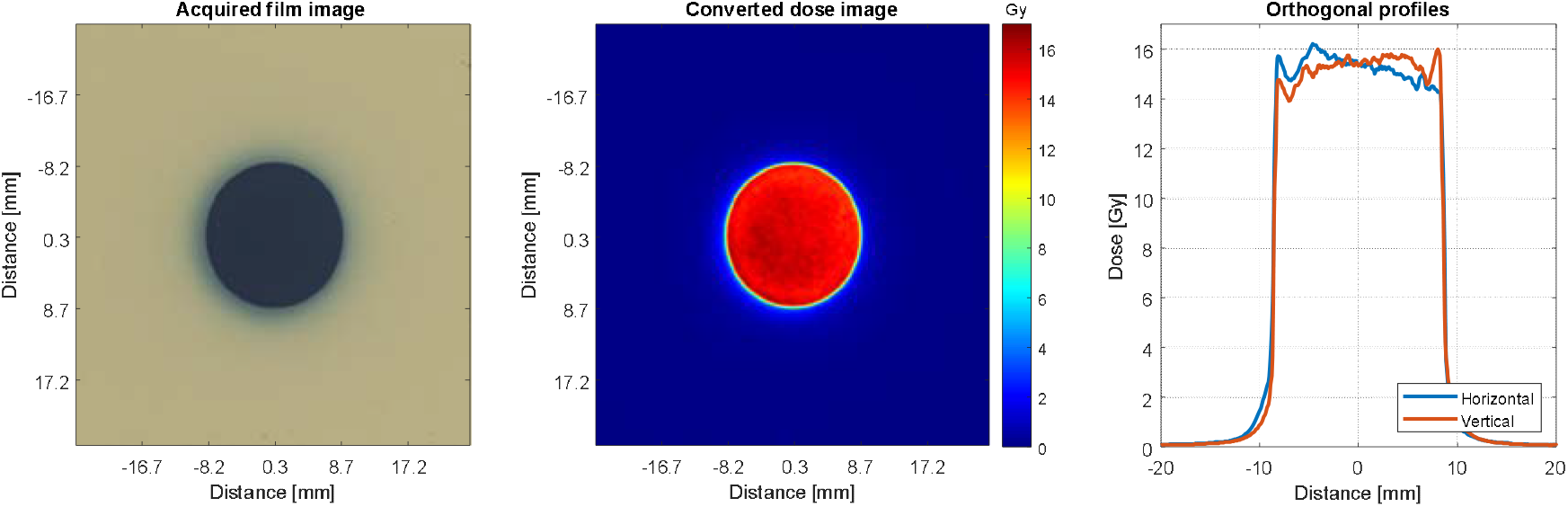
EBT3 measured dose at FLASH dose rate (110 Gy/s).

### 1.3 Cross-calibration of phantom detectors at PSI-Gantry 2

We have performed proton irradiations at PSI-Gantry 2 to check the reproducibility of phantom detectors and to determine correction factors for the specific beam quality by cross-calibration against a reference ion-chamber traceable to METAS.

The phantom detectors (Alanine, TLDs and EBT3 films) were loaded into a dedicated holder that matches the dimensions of an Advanced Markus (PTW, Freiburg, Germany) ion-chamber. The holder can allocate up to 7 Alanine/TLD pellets and 14 laser-cut EBT3 films (Figure 6). The cap of the holder has the same water equivalent thickness as the Advanced Markus build-up cap. 7 × 3 times Alanine pellets, 7 × 3 times TLD pellets and 14 × 2 times EBT3 films have been irradiated. Before that, the dose was measured 3 times with the Advanced Markus ion-chamber. All detectors were irradiated at isocenter at a water-equivalent depth of 12.7 mm. Beam parameters were as follows:

- Initial energy 170 MeV – same energy as for the biological experiments at Gantry1
- Beam spots arranged in a grid with equal spacing (2.5 mm) and weights.
- Field dimensions 10 × 10 cm^2^
- Field uniformity < 1%
- Field reproducibility < 0.1%

**Supplementary Figure 6.**
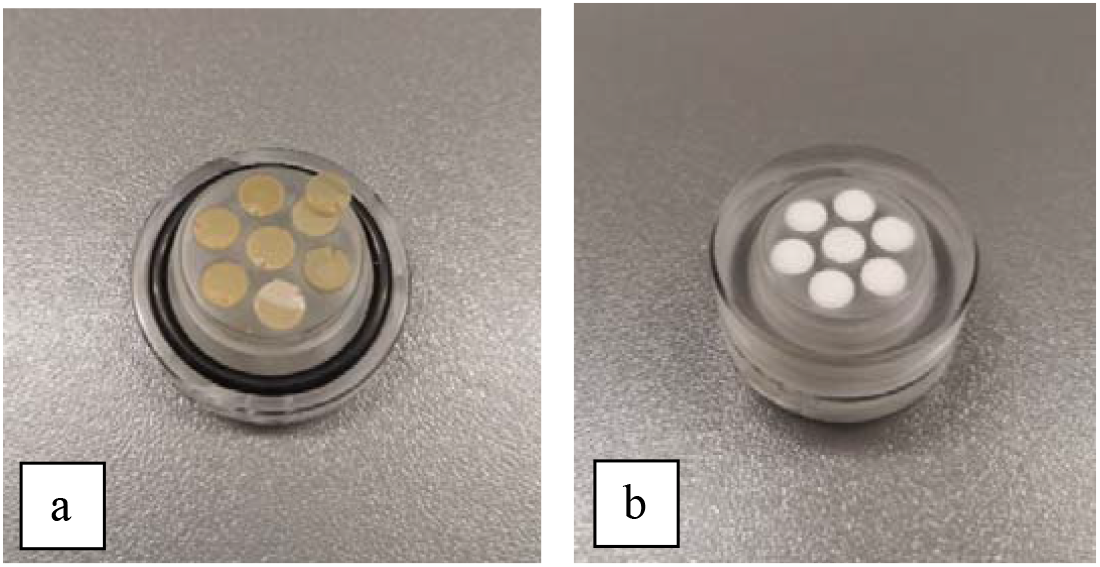
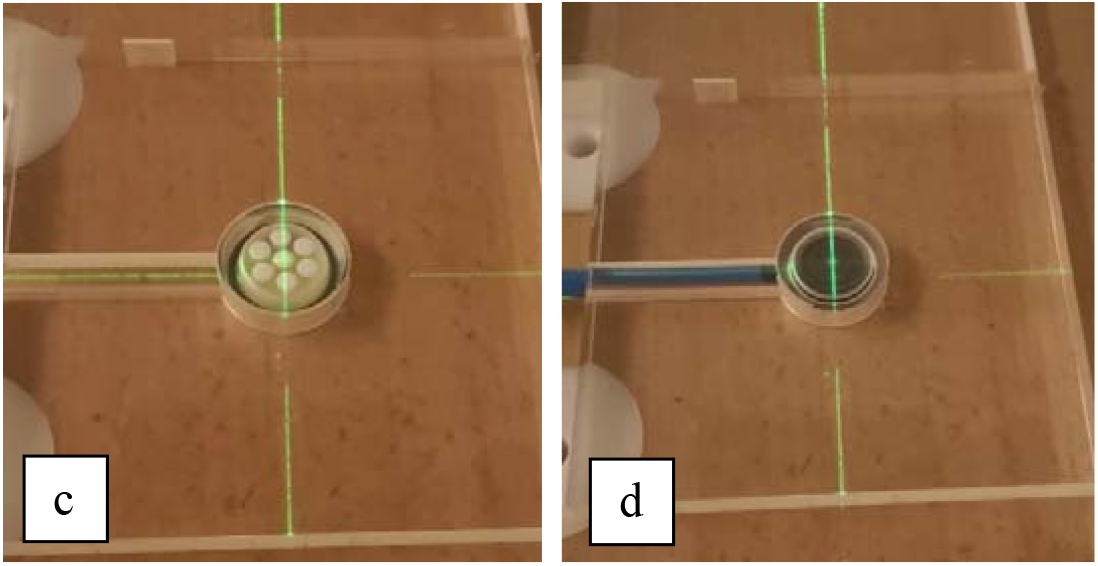
Images of the dosimeters loaded into the dedicated holder: a) laser-cut EBT3 films, b) and c) Alanine pellets. The Advanced Markus (d) chamber was used as a reference detector.

In Figure 7, the dose measured by the different detectors is plotted for all subsequent irradiations. For Alanine, TLDs and EBT3 films, each point represents the mean value of the readings within a single irradiation, and the error bars represent the standard deviation of the detectors loaded into the holder. The number of detectors irradiated per delivery is reported on the x-axis. Additionally, the mean values over multiple irradiations and the related standard deviation of the mean are shown for each detector type. For the Advanced Markus, only a single data point is shown, being its reproducibility <0.3 %.

The measured reproducibility of the phantom detectors was 0.9 %, 3.7 % and 1.4 % for Alanine, TLDs and EBT3 films, respectively.

**Supplementary Figure 7.**
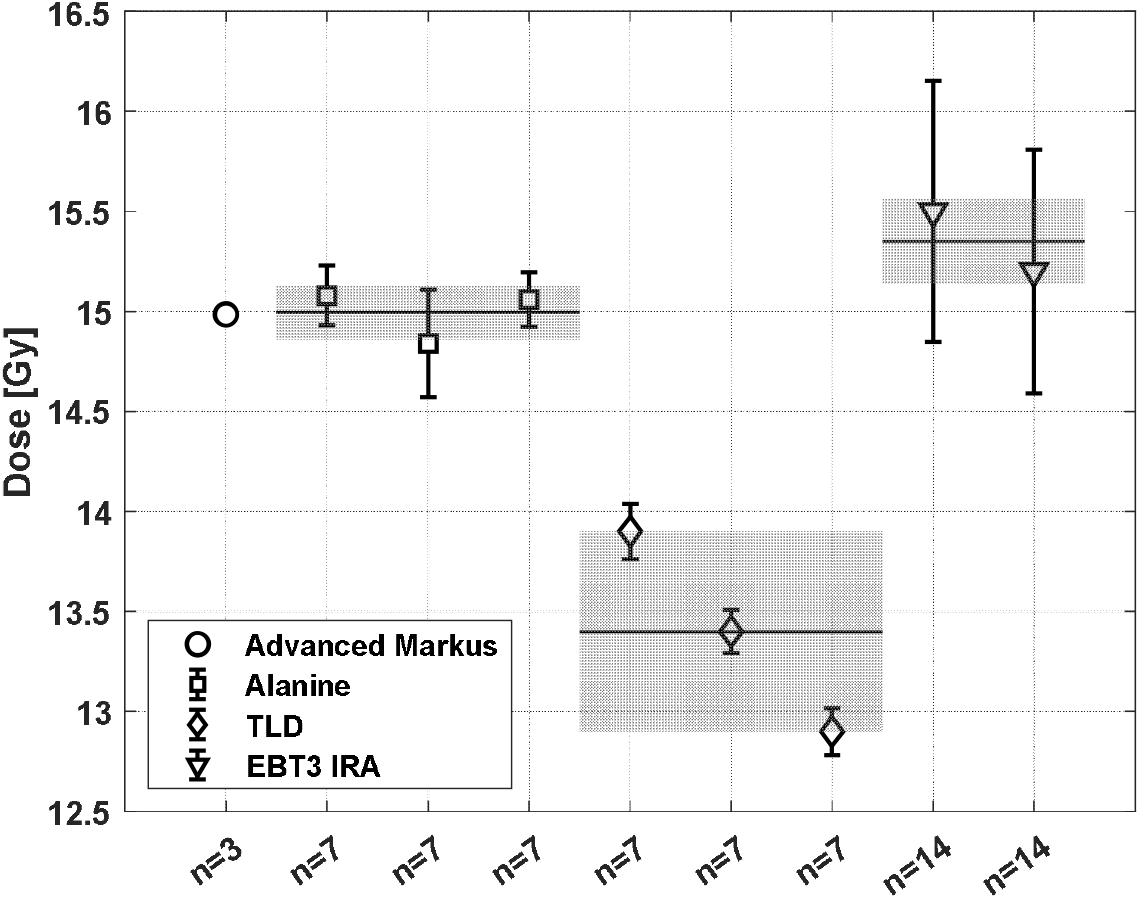
Measured dose for repeated irradiations at PSI-Gantry 2. The number of detectors irradiated in each delivery is reported on the x-axis. The error bars represent the spread of the readings in each delivery (the Advanced Markus bars are smaller than the marker size). The horizontal black lines are the mean values, and the shaded areas are the standard deviation of the mean values of multiple deliveries for each detector type.

The dose measured by the Advanced Markus in the proton single-energy field can be expressed using IAEA TRS-398 formalism:

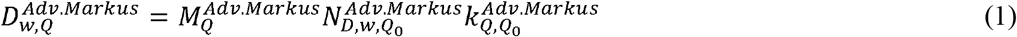

Where 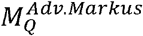 is the reading of the chamber corrected for ambient and chamber specific parameters, 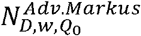 is the calibration factor determined at METAS in Co-60 reference beam, and 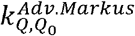 is the beam quality correction factor for the chamber at the beam quality *Q*. Similarly, the dose measured by each of the phantom detectors in the same proton field can be written as:

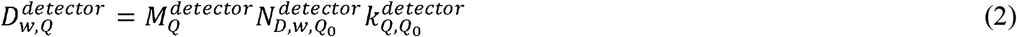

where the quantity 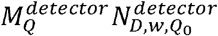 is provided directly by IRA after detectors readout and represents the absorbed dose to water in a Co-60 reference beam for the measured signal 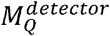.

Provided the irradiation field is the same for all detectors, it is possible to determine a beam quality correction factor for each of the phantom detectors by imposing

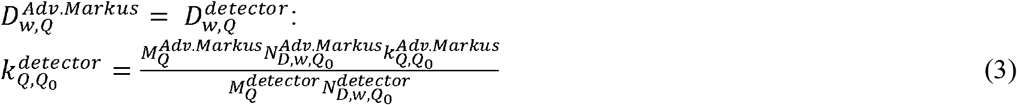

For this cross-calibration, we have chosen the same beam quality Q as used in the biological experiments at Gantry1, defined by an initial beam energy of 170 MeV and a measurement depth of 12.7 mm.

The beam quality correction factors were calculated according to equation (3) and found to be 1.00, 1.12 and 0.98 for Alanine, TLDs and EBT3 films, respectively. The associated uncertainties were estimated as in Palmans et al. study (including type-A uncertainties and uncertainties on 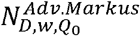 and 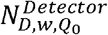) and amount to 2.3 %, 5.5 % and 2.5 %.

